# Negligible peptidome diversity of SARS-CoV-2 and its higher taxonomic ranks

**DOI:** 10.1101/2022.10.31.513750

**Authors:** Li Chuin Chong, Asif M. Khan

## Abstract

The unprecedented increase in SARS-CoV-2 sequence data limits the application of alignment-dependent approaches to study viral diversity. Herein, we applied our recently published UNIQmin, an alignment-free tool to study the protein sequence diversity of SARS-CoV-2 (sub-species) and its higher taxonomic lineage ranks (species, genus, and family). Only less than 0.5% of the reported SARS-CoV-2 protein sequences are required to represent the inherent viral peptidome diversity, which only increases to a mere ∼2% at the family rank. This is expected to remain relatively the same even with further increases in the sequence data. The findings have important implications in the design of vaccines, drugs, and diagnostics, whereby the number of sequences required for consideration of such studies is drastically reduced, short-circuiting the discovery process, while still providing for a systematic evaluation and coverage of the pathogen diversity.

## 1. Introduction

Sequence diversity is a major obstacle in the design of effective surveillance and intervention (vaccines, drugs, and diagnostics) strategies against viruses. Primary sequences are treasure troves for studies of sequence diversity, which can be alignment-dependent or alignment-free^1^. Big sequence data poses a major challenge to alignment-dependent approach, which is root to many sequence-based comparison studies. The approach is generally compute-intensive, time-consuming, and of reduced reliability with increase in number and variability of sequences, and thus, constraining its applications^1^. Multiple sequence alignment particularly becomes impractical when expanding the analysis to include all species under a genus, family or higher ranks of the taxonomic lineage. Towards this, an alignment-free approach offers an alternative strategy.

Our recently published tool, UNIQmin offers an alignment-free approach for the study of viral sequence diversity at any given rank of taxonomy lineage^1^. The tool performs an exhaustive search to generate a minimal set for a given sequence dataset of interest and is big data ready. The minimal set is the smallest possible number of distinct sequences required to represent a given peptidome diversity (pool of distinct peptides of a specific length, *k-*mer) exhibited by a dataset of interest, typically retrieved from public databases. This compression is possible through the removal of sequences that do not contribute effectively to the peptidome diversity pool, which is achieved by subjecting the retrieved sequence dataset to two levels of data compression. Firstly, through the removal of identical sequences, resulting in a deduplicated dataset, which is then followed by removal of unique sequences whose entire repertoire of overlapping *k-*mer(s) can be represented by other unique sequences in the dataset.

The utility of UNIQmin was demonstrated for the species *Dengue virus*, genus *Flavivirus*, family *Flaviviridae*, and the superkingdom *Viruses* (all datasets before the COVID-19 pandemic)^1^. Herein, we applied UNIQmin to protein sequence data of SARS-CoV-2 and its higher ranks of taxonomic lineages, namely species (with and without the SARS-CoV-2 sub-species), genus and family (**Figure 1**) to evaluate the effective viral sequence diversity at each rank^2^.

**Figure 1.**
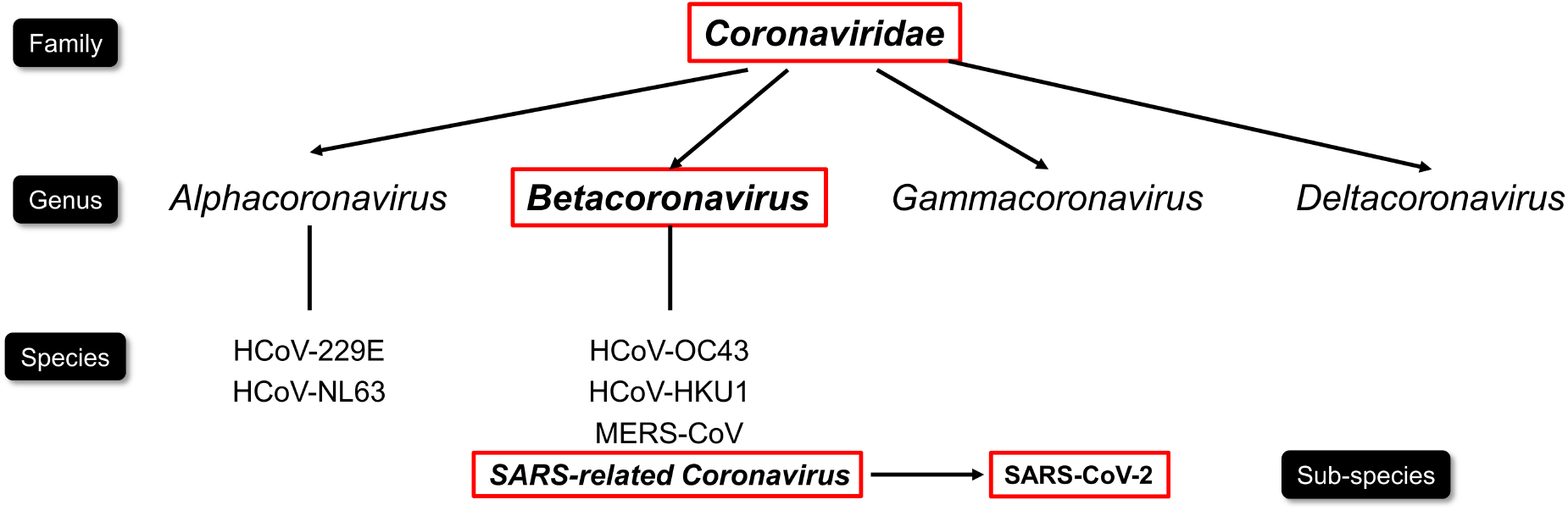
Taxonomic classification of coronaviruses known to infect humans (also known as HCoV).

## 2. Results

### 2.1. Extreme peptidome compression of SARS-CoV-2

A total of 56,340,320 (as of July 2021) protein sequence records of SARS-CoV-2 sub-species were retrieved from the GISAID EpiCoV™ database. Consequently, deduplication of the retrieved sequences for each of the 27 SARS-CoV-2 genome encoded proteins (**Figure 2(A)**) resulted in 1,780,901 unique sequences in total (∼3.2% of the retrieved dataset) (**Figure 2(B)**). Remarkably, this showcases that nearly all (∼96.8%) of the SARS-CoV-2 retrieved protein sequence records were identical. The same was observed for each of the 27 encoded, individual proteins, except the non-structural protein 3 (NSP3) and spike glycoprotein (S), both of which exhibited the largest fraction of unique sequences (∼16.6% and ∼16.9%, respectively) amongst the proteins. The high duplication of the virus protein records could be generally an outcome of spatio-temporal submission of identical/related strains, where sequence changes, if any, within a variant strain would be limited to one or more proteins, leaving the others unaltered. S protein (approx. 1,273 amino acids long), key for viral entry, exhibits multiple regions of high diversity (https://nextstrain.org/ncov/gisaid/global), with substitutions linked to the major variants identified thus far^3^, including the Omicron. This is in congruence with the lower duplication observed for S protein. In contrast, the NSP3 protein, which plays many roles in the viral life cycle, is generally of higher conservation than the S protein, with relatively fewer regions of high diversity. However, this largest protein of coronaviruses (approx. 7,096 amino acids long) is littered with numerous positions of low substitutions and is also reported to exhibit the highest number of missense mutations (547), followed by S (394)^4^. The total number of distinct substitutions across the length of different variants of the protein can affect duplicate removal. This may explain the lower duplication reduction for NSP3, which is comparable to S.

**Figure 2.**
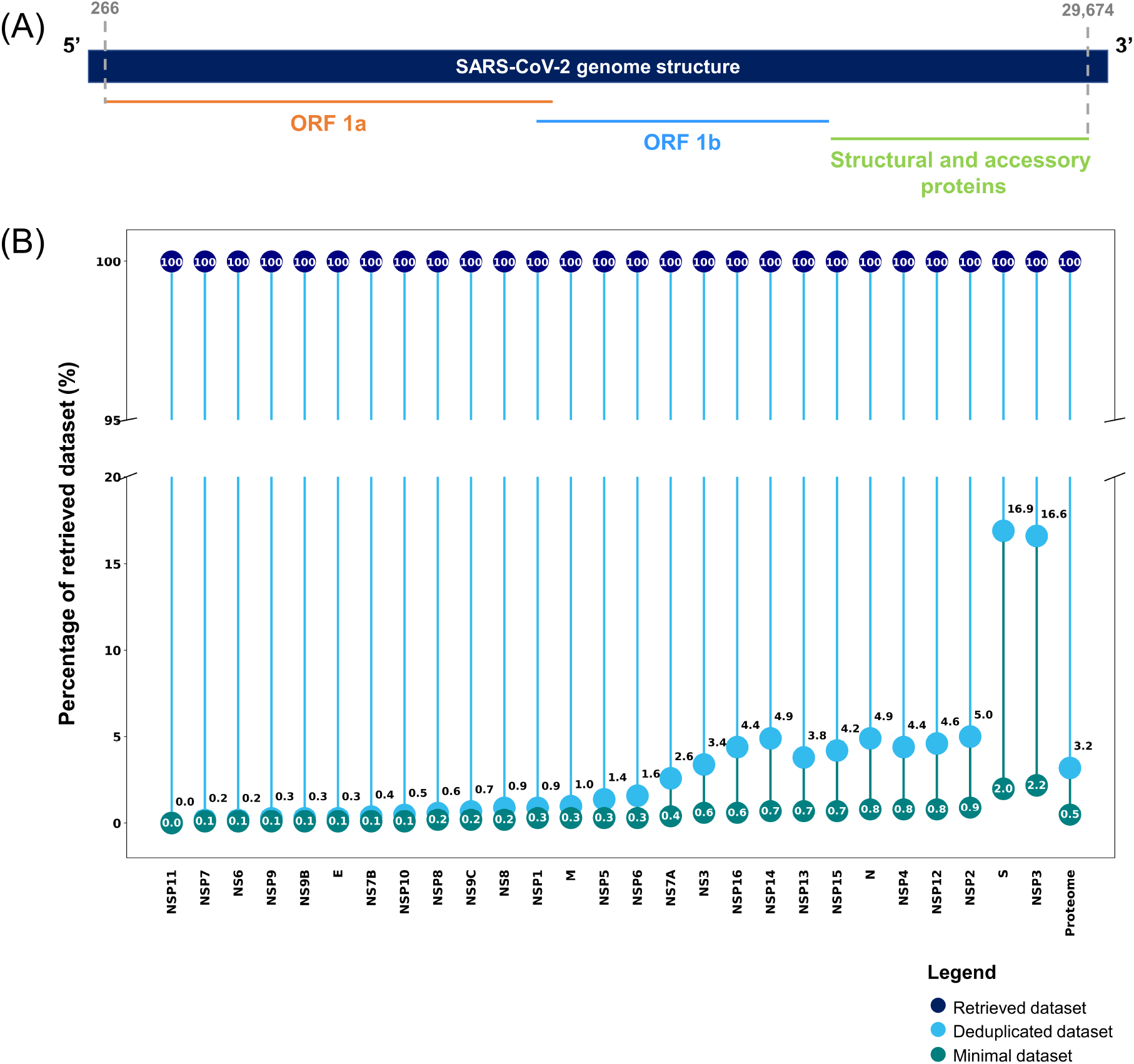
Compression of SARS-CoV-2 datasets across sub-species taxonomy lineage rank (proteins). (A) The genomic architecture of SARS-CoV-2. The genome encodes two large genes, ORF1a (orange) and ORF1b (blue) that collectively encode 16 non-structural proteins (NSP1-16), besides the four structural genes (spike (S), envelope (E), membrane (M), and nucleocapsid (N)), and five accessory proteins. (B) Compression for each protein of SARS-CoV-2. For instance, spike glycoprotein (S) had 2,115,156 sequences (100%; dark blue circle) in the retrieved dataset and compression resulted in a deduplicated dataset of 358,096 sequences (∼16.9% of the retrieved data; sky blue circle) and a minimal dataset of 42,399 sequences (∼2.0% of the retrieved data; teal circle). The percentages of the retrieved dataset are rounded to one decimal place; “0.0” for NSP11 is due to rounding and is actually < 0.1.

The 27 protein deduplicated datasets were then further compressed, through the removal of unique sequences by use of UNIQmin without incurring any loss of information in terms of the total peptidome repertoire (relevant to the *k*-mer of choice). The overlapping *k*-mer size of nine (9-mer or nonamer) was selected for immunological applications (see Chong et al., 2021 for various considerations on *k*-mer size). The resulting 27 protein minimal datasets totalled 273,851 sequences (∼0.5% of the retrieved dataset) (**Figure 2(B);** minimal datasets are available at https://github.com/ChongLC/UNIQmin_PublicationData/tree/main/ApplicationPaper_SARS-CoV-2). Strikingly, only less than 0.5% (a total reduction of ∼99.5%) of the more than 56 million reported SARS-CoV-2 protein sequences are required to represent the inherent viral peptidome diversity. The minimal sets were uniformly small for each of the proteins, including for S and NSP3, though they were relatively higher for these two (∼2.0% to ∼2.3%, respectively) compared to the other proteins (<1%).

### 2.2. All non-SARS-CoV-2 sub-species are more diverse

All reported viral protein sequences for the species *SARS-related coronavirus*, without SARS-CoV-2 sub-species, were retrieved from the NCBI Entrez Protein database (as of July 2021) for comparative analysis. The retrieved dataset of 4,612 sequences was deduplicated and removed of unique sequences that did not effectively contribute to the peptidome diversity (**Figure 3**). This resulted in a deduplicated dataset of 938 sequences (∼20.3% of the retrieved data; reduction of ∼79.7%) and a minimal dataset of 681 sequences (∼14.8% of the retrieved data; total reduction of ∼85.2%). The species *SARS-related coronavirus*, excluding SARS-CoV-2, comprised of sequences that are much more diverse than those of the COVID-19 disease agent.

**Figure 3.**
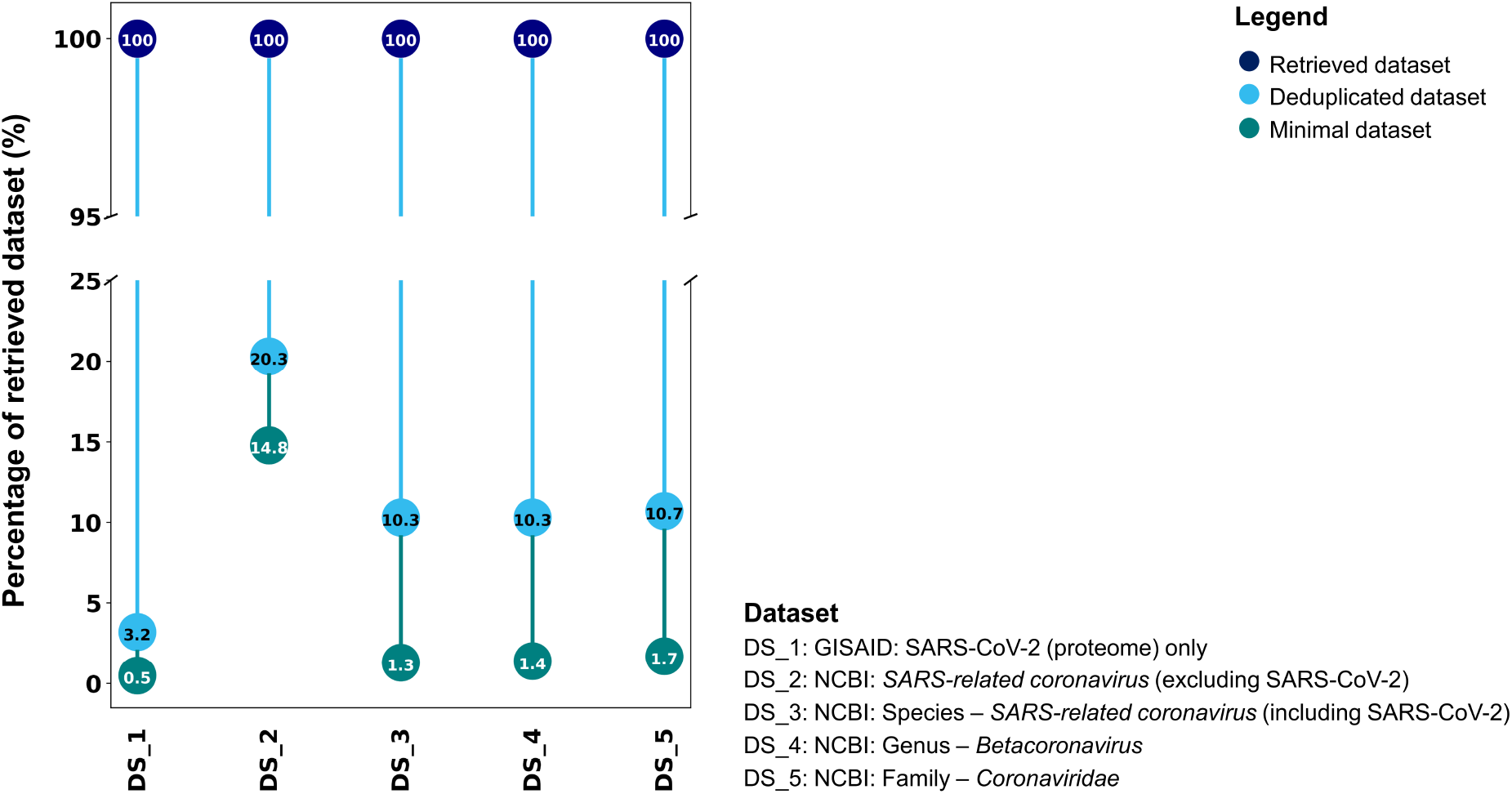
Compression for all retrieved protein datasets across SARS-CoV-2 taxonomic lineage ranks, namely sub-species, species (with and without SARS-CoV-2), genus, and family.

### 2.3. Linear compression trend at species, genus and family ranks

The diversity analysis was further expanded to higher ranks of the SARS-CoV-2 taxonomy lineage (https://www.ncbi.nlm.nih.gov/taxonomy), namely species *SARS-related coronavirus* (including SARS-CoV-2), genus *Betacoronavirus*, and family *Coronaviridae*. This analysis could not be done by use of the data available at GISAID as the database only provides the SARS-CoV-2 dataset, and thus, the NCBI Entrez Protein database was used. Protein sequences retrieved (as of July 2021) for the species, genus and family totalled 4,669,400, 4,689,400, and 4,733,200 records, respectively. The genus *Betacoronavirus* had an additional of only 20,000 sequences representing all the 213 species within the genus, besides the *SARS-related coronavirus*. This number more than doubled to 43,800 as additional sequences representing all the other genera under the family *Coronaviridae*. A summary of the compressions is shown in **Figure 3**. Deduplication of the three datasets resulted in near-identical reductions, ranging from ∼89.3% (∼10.7% of the retrieved family dataset) to ∼89.7% (∼10.3% of the retrieved species and genus datasets), indicating that the main duplicate reduction was at the species rank (contributed largely by SARS-CoV-2), after which there was near-no change at the genus rank and only a negligible at the family rank. This underscores that the other species within the genus *Betacoronavirus*, which includes notable human pathogens (such as SARS-CoV, MERS-CoV, and seasonal HCoV-HKU1 and HCoV-OC43), and the species of the other genera of the family *Coronaviridae* (includes human pathogens, seasonal HCoV-229E and HCoV-NL63)^5^, are understudied. Each deduplicated dataset was further compressed of unique sequences, resulting in minimal datasets of 61,819 (∼1.3% of the retrieved species data), 65,117 (∼1.3% of the retrieved genus data), and 79,414 (∼1.7% of the retrieved family data) sequences, respectively.

The significantly reduced minimal dataset of the genera (∼98.7%) and family (∼98.3%) suggested that only less than 2% of the sequences are alone sufficient to capture the inherent diversity. This highlights that an additional of 3,298 sequences were sufficient to capture the diversity of all other species of the genus *Betacoronavirus*, besides *SARS-related coronavirus*. A further 14,297 sequences were sufficient to represent all species of the other genera under the family *Coronaviridae*.

## 3. Conclusions

Just within two years of the pandemic, the unprecedented growth of SARS-CoV-2 sequence data has become a benchmark for sequencing and studies of other notable pathogens. The data tsunami has strained and challenged the current bioinformatics tools, pipelines, and methodologies for handling and mining of big data. For example, the recently released multiple sequence alignment tool, MAGUS^6^ are only able to effectively align several hundred thousand sequences reliably and within a reasonable time, when there are more than 13 million protein sequences publicly available (as of October 2022) for analysis just for the spike protein alone. The alignment-free approach described herein is applicable to both DNA and protein sequences and mitigates the need for alignment to study diversity. Moreover, the data compression through the generation of a minimal set can potentially, in turn, enable application of alignment approaches to the much-reduced dataset. More importantly, the key finding herein is that less than 0.5% of the ∼56 million protein sequences of SARS-CoV-2 are alone sufficient to represent the inherent peptidome diversity, which only increases to a mere ∼2% at the family lineage rank. These percentages of the minimal set are expected to remain relatively the same even with further increases in the sequence data. This is because as data grows, viral variants exhibiting novel *k*-mers relative to the existing repertoire of sequence diversity are likely to diminish, particularly for the SARS-CoV-2 sub-species given the already large dataset, or show limited growth at the genus and family ranks; unless, the existing repertoire is perturbed by discovery of sequences of previously unknown members. The results herein have important implications to effective viral surveillance strategies and vaccine, drug, and diagnostic designs, whereby the number of sequences required for consideration of such studies can be drastically reduced, short-circuiting the discovery process, while still providing for a systematic evaluation and coverage of the pathogen diversity.

## 4. Methods

### Sequence data retrieval and pre-processing

All reported protein sequences of SARS-CoV-2 were downloaded from the GISAID EpiCoV™ database (https://www.gisaid.org; as of July 2021)^7^ and were subsequently separated into the individual 27 encoded proteins based on the annotations in the sequence header (description) line of the retrieved FASTA file. Additionally, all reported protein sequences of the species *SARS-related coronavirus* (taxonomy ID: 694009), genus *Betacoronavirus* (taxonomy ID: 694002), and family *Coronaviridae* (taxonomy ID: 11118) were retrieved from the NCBI Entrez Protein Database^8^ using NCBI Mass Sequence Downloader^9^ (as of July 2021). In order to also analyse the species dataset without the SARS-CoV-2 sub-species, all reported sequences of SARS-CoV-2 (taxonomy ID: 2697049) were filtered out with the help of the sequence header annotation, facilitated by the tool, faSomeRecords (https://github.com/santiagosnchez/faSomeRecords).

### Sequence deduplication and generation of minimal sets

The clustering tool, CD-HIT^10^ was used to remove sequence duplicates from each of the database retrieved datasets. The tool, UNIQmin^1^ was used to generate a minimal set for each of the deduplicated datasets.

## Data Availability

The datasets were derived from sources in the public domain, namely GISAID EpiCoV™ and NCBI Entrez Protein databases, while the data underlying this article is available at GitHub: https://github.com/ChongLC/UNIQmin_PublicationData/tree/main/ApplicationPaper_SARS-CoV-2.

## Author Contributions

AMK conceived the initial idea, supervised the analysis, reviewed/amended the manuscript. LCC contributed to data curation, analysis, visualisation, writing of the original manuscript draft, and made the necessary improvements upon the reviews. Both authors approved the final version.

## Acknowledgments

We gratefully acknowledge the authors from the originating and submitting laboratories for the sequences deposited to GISAID’s EpiCoV™ database and also NCBI Protein database.

